# The human hippocampus is involved in auditory short-term memory trace formation

**DOI:** 10.64898/2025.12.01.691561

**Authors:** Xiuyuan Liang, Zihao Guo, Dong Zhang, Yiming Zhang, Aobo Chen, Lanlan Wang, Xiang Liu, Wang Zhang, Yinbao Qi, Xiaorui Fei, Ming Wang, Huawei Li, Lin Chen, Ruobing Qian

## Abstract

The human hippocampus has been claimed to play an important role in long-term memory, or episodic memory, but its contribution to short-term memory remains debated. Here, we demonstrate human hippocampal involvement in auditory short-term memory (ASTM) trace formation. We used a classic oddball paradigm and intracranial recordings across various human brain areas. High frequency activities evoked by deviant stimuli in subjects indicate the generation of ASTM trace. Analysis of auditory response latencies across the hippocampus, insula, and temporal, parietal, and frontal lobes revealed the early processing of ASTM trace at a pre-attention stage. Granger causality analysis further showed that ASTM trace processing engages hierarchical cortical areas and clarified their interactions. Bottom-up signals flow from the insula and auditory temporal regions to the frontal lobe early on, followed by top-down information from the frontal lobe to the hippocampus, inferior temporal lobe, and insula at a later pre-attentive stage. Critically, Bayesian model selection of dynamic causal modeling revealed that deviant stimuli selectively enhanced the effectiveness of top-down connections to the hippocampus, thereby facilitating ASTM trace formation. These results provide compelling evidence that the human hippocampus contributes to auditory short-term memory at a pre-attention stage and suggests a challenge to entrenched beliefs in the classification of memory systems.

## Introduction

The human hippocampus has been claimed to play a crucial role in human memory system especially long-term memory, also referred to as episodic memory^1^, it comes with an equally important caveat that the hippocampus does not contribute to short-term memory^2^. The traditional/conventional view of hippocampal amnesia is that short-term memory is preserved despite the severe impairment in long-term memory^1,3,4^. For several years, a wide range of studies have established the medial temporal lobe, and hippocampus in particular, to be critically associated with long-term memory processing. For instance, researchers studied H.M., the well-known amnesic patient who had undergone temporal lobe resection, and published observations of some types of preserved memory (motor skill and perceptual learning) in the patient^5,6^, as well as types of memory that depended on parts of the brain other than the medial temporal lobe, in particular the hippocampus, or that did not depend on conscious awareness were termed implicit (non-declarative) memory^7,8^. Previous view mentioned that amnesia associated with hippocampal region damage is selective to declarative memory, the ability to recall facts and events^9^, and for H.M. the deficit includes the complete absence of episodic memory, defined as the ability to remember specific personal experiences^10^. In addition, many researchers have explored the underlying neural mechanism of memory in which studies have implied that selective injury to the medial temporal lobe leads to an isolated deficit in long-term memory^11,12^. Evidence from studies in patients with amnesia due to hippocampal damage proposed that patients exhibit impairment of conscious episodic encoding and retrieval^13^. Nevertheless, there is different view reports that amnesic patients with bilateral lesions involving the hippocampus region suffered from an impairment of short-term memory along with research develops^14,15^. Moreover, recent visual study in human supports the view that the hippocampus does indeed have a role to play in short-term memory^16^ and study reported the role of cross-frequency coupling in cortico-hippocampal activity in sequential auditory short-term memory (ASTM)^17^.

Short-term memory describes the mental capacity to consciously hold information, such as keeping a number in mind. Its neural substrate is the memory trace. For example, the active representation of that number, through specific neuronal firing patterns in networks involving the hippocampus and prefrontal cortex, constitutes the short-term memory trace. Thus, the trace is the persistent neural state that enables the transient cognitive experience of short-term memory. Short-term memory is a storage system that has an evolutionary survival advantage, such as paying attention to limited but essential information excluding confounding factors and ASTM is one of the important sensory-associated subsystems^18^. Short-term memory trace is the physical neural basis for that function, the temporary, biological activity in our brain that makes short-term memory possible. Mismatch negativity (MMN), first described about 40 years ago by Näätänen and his colleagues^19^, is an index of automatic deviance detection. It reflects cortical responses to deviant changes in the auditory system^20,21^ and has been regarded as a marker for detecting a deviant stimulus within the ASTM^22^. A variety of studies have led to the understanding that the MMN is the outcome of an automatic comparison process between a new, deviant stimulus and the memory trace formed by the sensory representation of the standard stimuli within the short-term memory^23–26^. The MMN depends on the presence of ASTM trace in the auditory cortex representing the repetitive aspects of the preceding auditory events^20^. Thus, ASTM trace formation is indexed by the MMN responses and deviance detection is mediated by a more stable short-term memory trace^27,28^.

In the current study, we employed intracranial recordings, a technique affording high spatial and temporal resolution, across various human brain regions, combined with a short-term memory paradigm presented with a contrast pure tone auditory streams to explore whether the human hippocampus contributes to ASTM trace formation. High frequency activities (HFA, 70–150 Hz) elicited by the auditory stimuli is a reliable electrophysiological correlate of underlying averaged spiking activity generated by the thousands of neurons that are in the immediate vicinity of the recording electrodes^29–32^. This type of activity has been linked to various cognitive processes, including attention, memory, and sensory processing. Significant HFA of deviance detection evoked by the subjects indicate the generation of ASTM trace. In our study, HFA results revealed that hippocampus was involved during process of ASTM trace. Auditory response latencies recorded from the hippocampus, insula, temporal lobe, parietal lobe, and frontal lobe showed the early processing of ASTM trace at a pre-attention stage. Moreover, Granger causality analysis further demonstrated ASTM trace is processed in hierarchical cortical areas, and detailed interactions of brain regions among this memory trace process have been clarified/interpreted clearly. Specifically, bottom-up auditory signals processed by the insula, and auditory regions of the temporal lobe and the frontal lobe at an early stage, then auditory top-down information transmitted from the frontal lobe to the hippocampus, the inferior temporal lobe, as well as insula at a relatively late pre-attention stage of the auditory information processing. Although Granger causality characterizes the temporal precedence and statistical dependencies of directional information flow, it does not specify the underlying neurobiological mechanism. To identify the specific neural mechanism, we employed Dynamic Causal Modeling (DCM). Critically, DCM results revealed that deviant stimuli selectively enhance top-down connections from the frontal, insula or temporal to the hippocampus, providing a mechanistic explanation for the information flow that facilitates ASTM trace formation. In summary, our findings provide converging evidence that the human hippocampus contributes to ASTM trace formation at a pre-attentive stage through a specific frontal-hippocampal neural circuit that underlies the formation of this transient memory trace, suggesting a new perspective that challenges traditional classifications of memory systems.

## Results

### High frequency responses to auditory deviants across hippocampal and cortical networks

To investigate the neural mechanisms underlying short-term memory trace formation, we employed a classic auditory oddball paradigm combined with intracranial recordings across multiple human brain regions. The hippocampus, a key structure within the medial temporal lobe memory system, is critically involved in the integration and relay of auditory information to downstream perceptual and cognitive regions^33–36^. Thus, for each recording channel in the hippocampal region, we computed trial-averaged high-frequency activity responses to both standard and deviant pure tones (Fig. 2). Our results revealed that hippocampal channels displayed diverse response profiles, including both HFA enhancements and high-frequency suppression (HFS). Specifically, responses to standard stimuli were predominantly characterized by negative-going deflections (Fig. 2a, c), whereas responses to deviant stimuli consistently exhibited positive deflections (Fig. 2e, g). To identify channels exhibiting significant responses to auditory stimuli, a two-tailed cluster-based permutation test incorporating threshold-free cluster enhancement (TFCE)^37^ was applied separately to standard and deviant trials for each channel. The test was performed using 1000 sign-flipping randomizations. Channels with a corrected p-value below 0.05 for either stimulus type were classified as responsive. Exemplar channels exhibiting statistically significant responses to standard pure tones are shown in or deviant pure tones are shown in Fig. 2b and Fig. 2d (*p* < 0.05), while exemplar channels exhibiting statistically significant responses to deviant pure tones are presented in Fig. 2f and Fig. 2h (*p* < 0.05).

**Figure 1.**
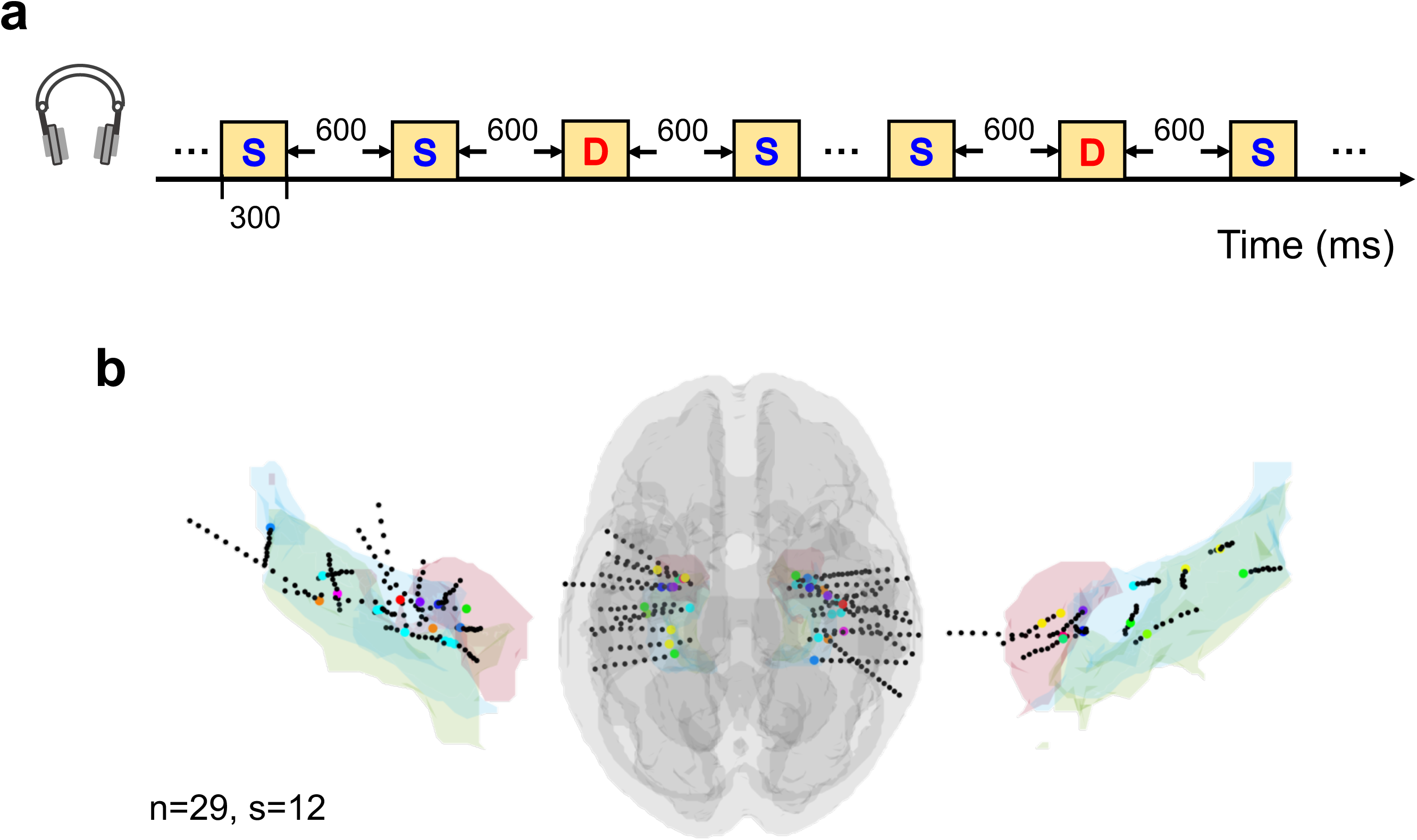

**Figure 2.**
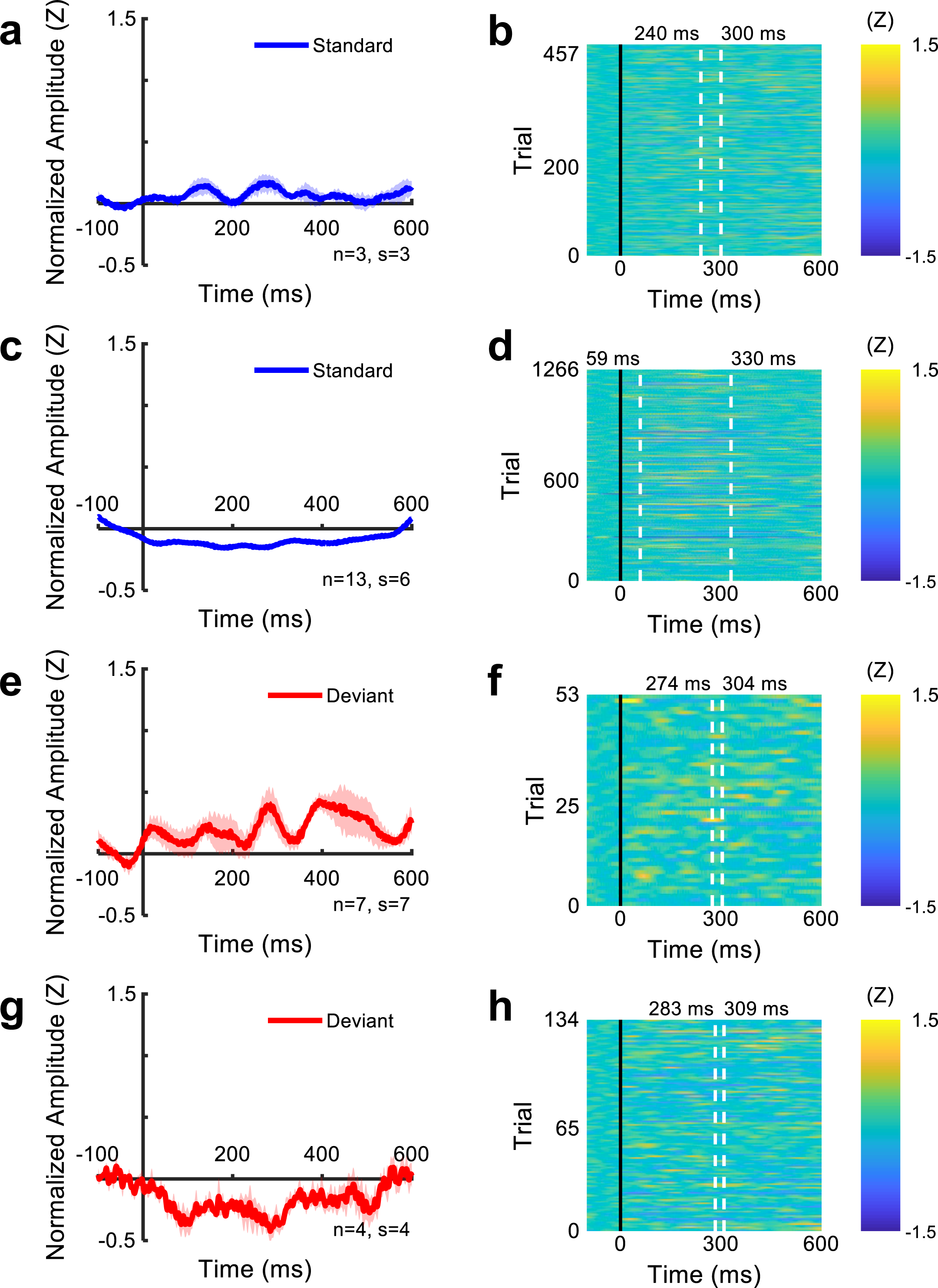

Furthermore, electrode pairs that evoked significant mismatched activity were categorized according to whether the mismatch activity was enhanced or suppressed. We collected sEEG data from 17 epilepsy patients (Table 1). To obtain a global description of the brain regions in deviance detection, we examined the HFA responses to standard versus deviant tones across intracranial channels. In the HFA analysis, 49 channels responded to tone responses in 10 out of the 17 patients (Fig. 3), which illustrates the overall response profiles across all channels, including both HFA and high-frequency suppression (HFS). Specifically, Significant HFS responses were observed in 8 channels across 6 patients with and 41 channels across 10 subjects with HFA responses (*p* < 0.05, Fig. 3a, b). HFS in response to the stimuli at a latency of 395 ms were recognized at the middle temporal gyrus (MTG), temporo-occipital brain area, posterior inferior temporal gyrus, and anterior corona radiata a channel. In addition, 29 channels across 9 patients responded exclusively to deviant stimuli (Fig. 3c) whereas 12 channels across 7 patients responded to both standard and deviant tones (Fig. 3d). Supplementary Figure 1 showed HFA responses across individual subjects. It is known that studying HFA with sEEG recordings can provide insights into the neural bases of human cognition that cannot be derived as easily from the study of BOLD effects with fMRI or from the analysis of EEG^30^. HFS, a transient energy suppression in the gamma-band (40−150 Hz) during the processing of sensory stimuli, which was first reported by Lachaux^38^ and has close relationship with attention^39^. We also observed widely distributed high frequency suppression (HFS)^40^, particularly in response to standard stimuli, which generally occurred later than high frequency activity (HFA). This widespread HFS may reflect an energy saving mechanism in the brain when processing predictable events. Given the known interaction between early tone responses and later repetition suppression^41^, such suppression may be mediated by repetition suppression mechanisms.

**Figure 3.**
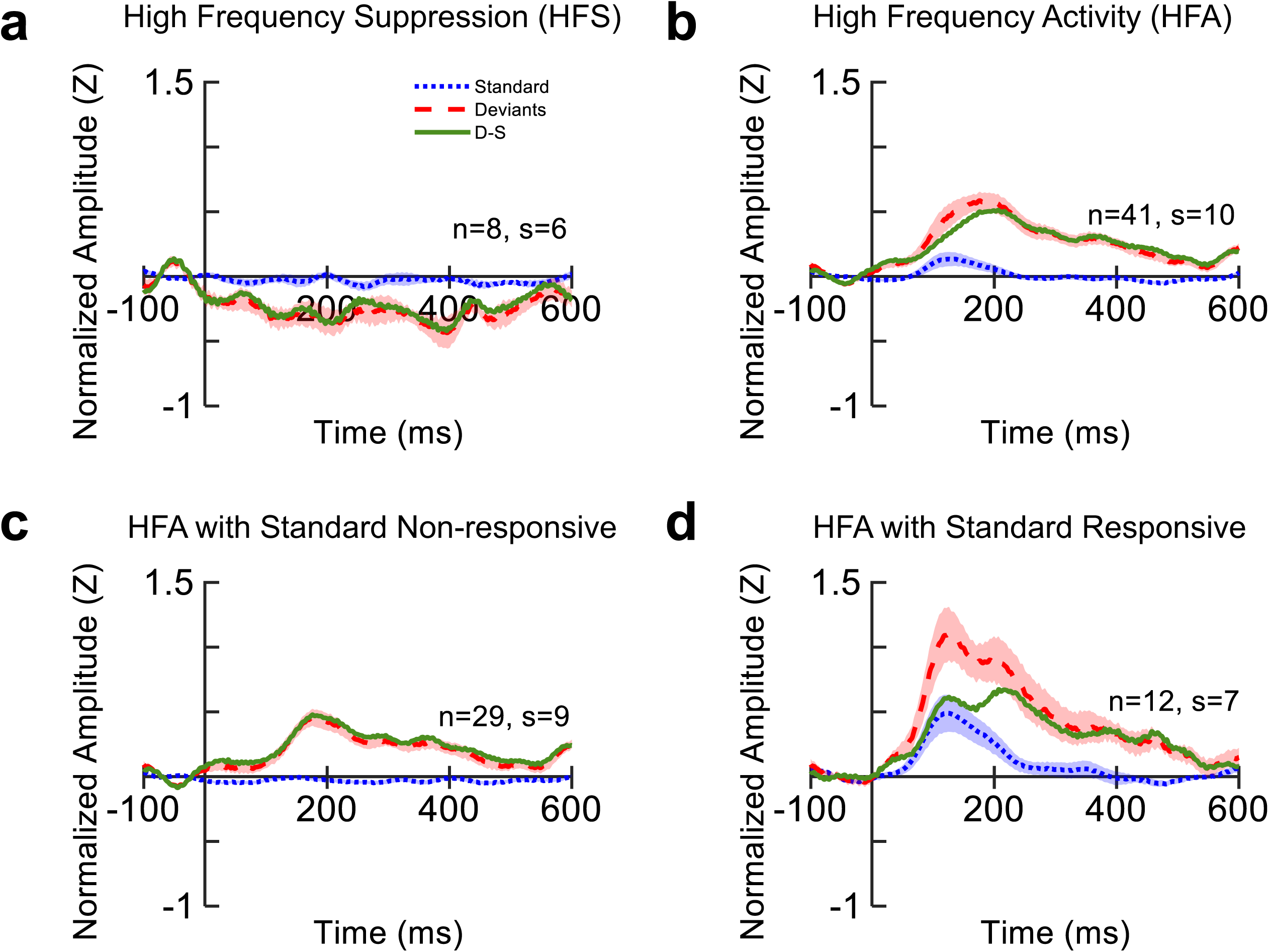

### Spatiotemporal responses of deviance processing

Auditory deviance elicited HFA responses across a distributed cortical network, with response latencies following a systematic spatial order in individual subjects (Fig. S1). Mean HFA/HFS response waveforms for key brain regions to standard and deviant pure tones, along with the deviant-minus-standard difference waveforms were shown for multiple brain regions (Fig. 4a). Brain regions were defined as follows: the insula; temporal lobe (encompassing Heschl’s gyrus, superior temporal gyrus, and superior temporal sulcus); frontal lobe (including the middle frontal gyrus, superior frontal gyrus, anterior cingulate cortex, and frontal operculum); parietal lobe (comprising the postcentral gyrus, parietal operculum, and supramarginal gyrus); posterior temporal lobe (the temporo-occipital part of the middle temporal gyrus); inferior temporal lobe (including the inferior temporal gyrus, fusiform gyrus, and inferior portion of the middle temporal gyrus); and the hippocampal area (encompassing the hippocampus, parahippocampal gyrus, and amygdala). Additionally, analysis of response latencies across these regions revealed a precise spatiotemporal gradient (Fig. 4b). The earliest responses occurred in the insula and primary auditory cortex (∼110 ms), followed by sequential engagement of the frontal and parietal cortices. The latest responses were observed in the hippocampus and inferior temporal cortex, with latencies extending up to ∼438 ms. This ascending latency pattern demonstrates that deviance processing propagates from early sensory to higher order associative regions.

**Figure 4.**
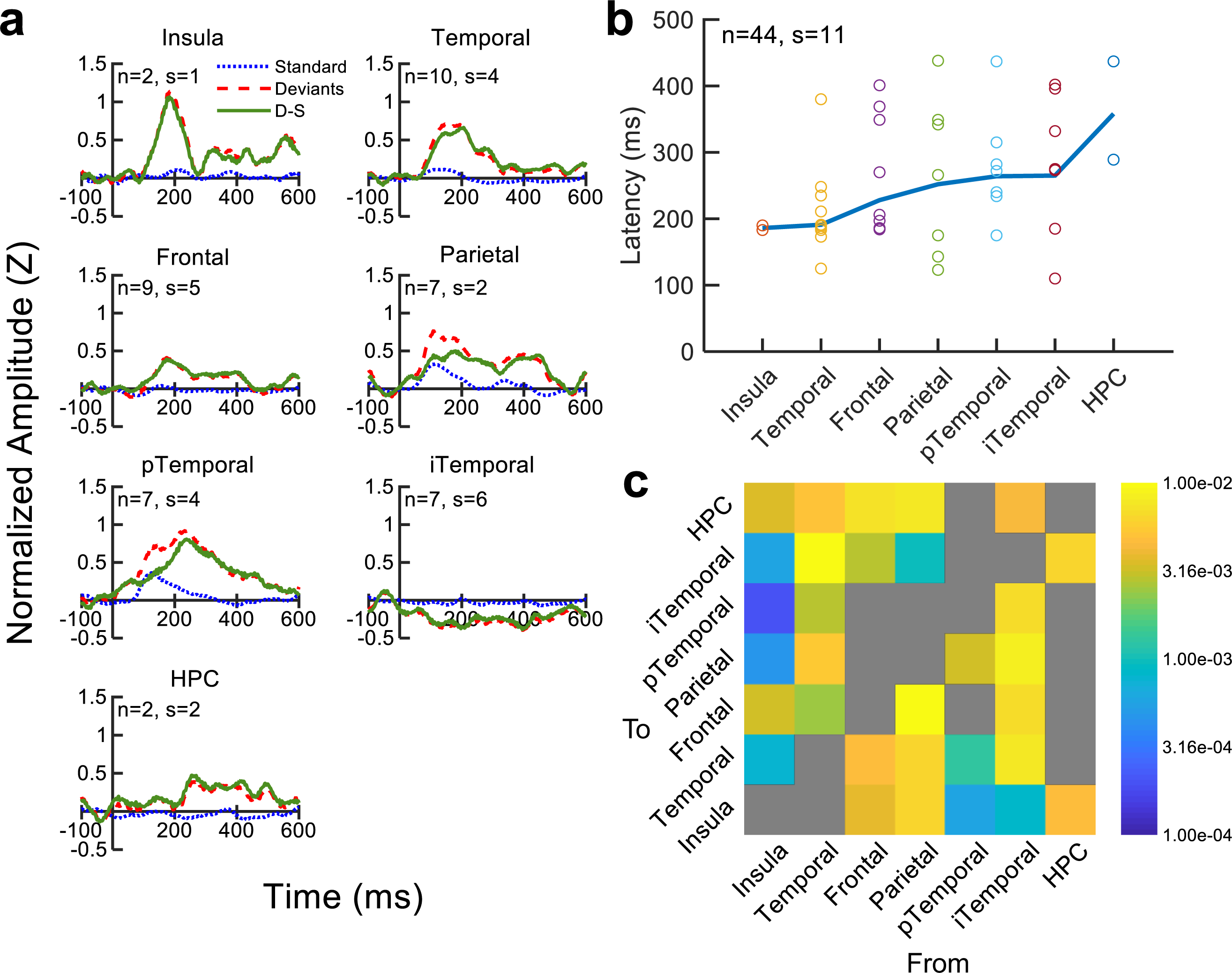

### Granger causality analysis revealed hierarchical information flow between the hippocampus and various brain regions

To investigate the neural functional connectivity underlying auditory deviance processing, we applied linear Granger causality (GC) analysis. The multivariate Granger causality analysis (MVGC) Matlab toolbox used in the current study was designed for applications in neuroscience^42^, which implements numerical routines for calculating multivariate Granger causality (MVGC) from time series data, both unconditional and conditional, in the time and frequency domains^43^. The anatomical locations of these responsive brain regions (regions detailed above) were mapped onto a common cortical space, as shown in the enlarged image of the boxed area, with different colors denoting distinct brain clusters. The functional connectivity is depicted in the heatmap (Fig. 4c), which displays the estimated difference in Granger causal connection strengths between deviant and standard stimulus-evoked activities. This GC analysis revealed a bottom-up information flow from the insula and auditory-related regions to the frontal lobe, followed by the top-down information flow from the frontal lobe back to the hippocampus, inferior temporal lobe, and insula at the pre-attentive stage (Fig. 4c). This spatiotemporal profile establishes the hippocampus as a key node receiving feedback signals within the pre-attentive processing. A magnified inset of the demarcated area is presented, in which unique colors code for separate functional clusters and the directionality (bottom-up versus top-down) of their causal influences (Fig. 5).

**Figure 5.**
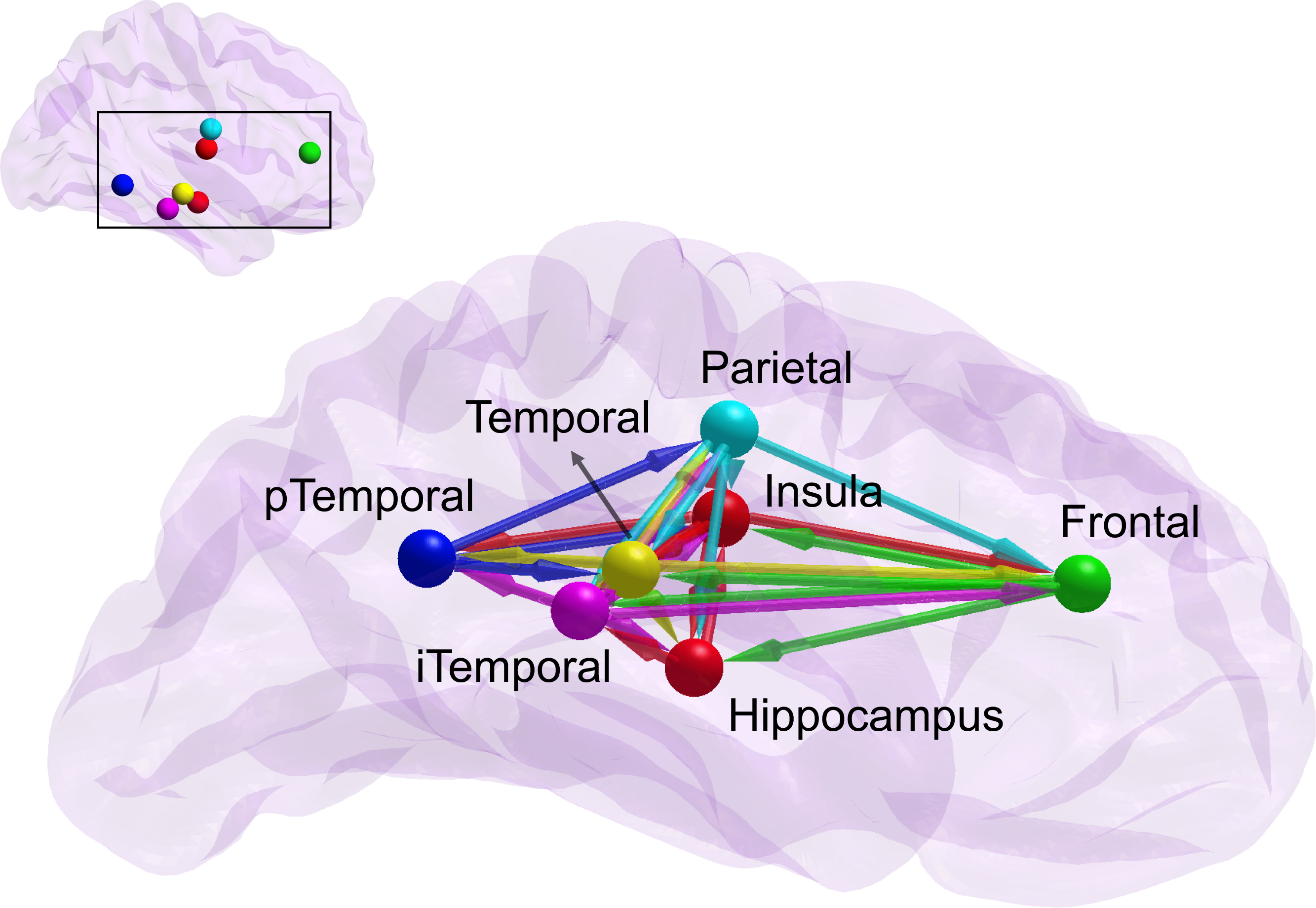

### Dynamic causal modeling revealed stimulus gated enhancement of top-down hippocampal connectivity

Bayesian Model Selection based on dynamic causal modeling yielded distinct winning models for each subject, indicating individual variability in the optimal circuit architecture. For Subject 3, Model 2 was identified as optimal (Fig. 6a, b). The key feature of this model is that it included a top-down connection from the temporal cortex to the hippocampus, which was significantly enhanced by deviant stimuli. For Subject 16, Model 7 was selected as the winning model (Fig. 6c, d). This model is characterized by the inclusion of top-down connections from both the frontal cortex and the insula to the hippocampus, with both connections showing significant enhancement during deviant processing. Bayesian parameter estimation of the respective winning models confirmed that deviant stimuli selectively and significantly enhanced the effectiveness of the identified top-down connections to the hippocampus in both subjects. In summary, the DCM analysis identified distinct and subject-specific optimal models supporting ASTM trace formation. Despite individual differences in the precise cortical nodes driving the hippocampus, the core mechanistic finding was consistent: deviant processing is characterized by a selective enhancement of top-down effective connectivity directed at the hippocampus. This result provides a mechanistic neural explanation for the Granger causality findings, demonstrating that the observed top-down information flow is supported by a dynamic, stimulus-dependent increase in synaptic efficacy within direct hippocampal afferent pathways.

**Figure 6.**
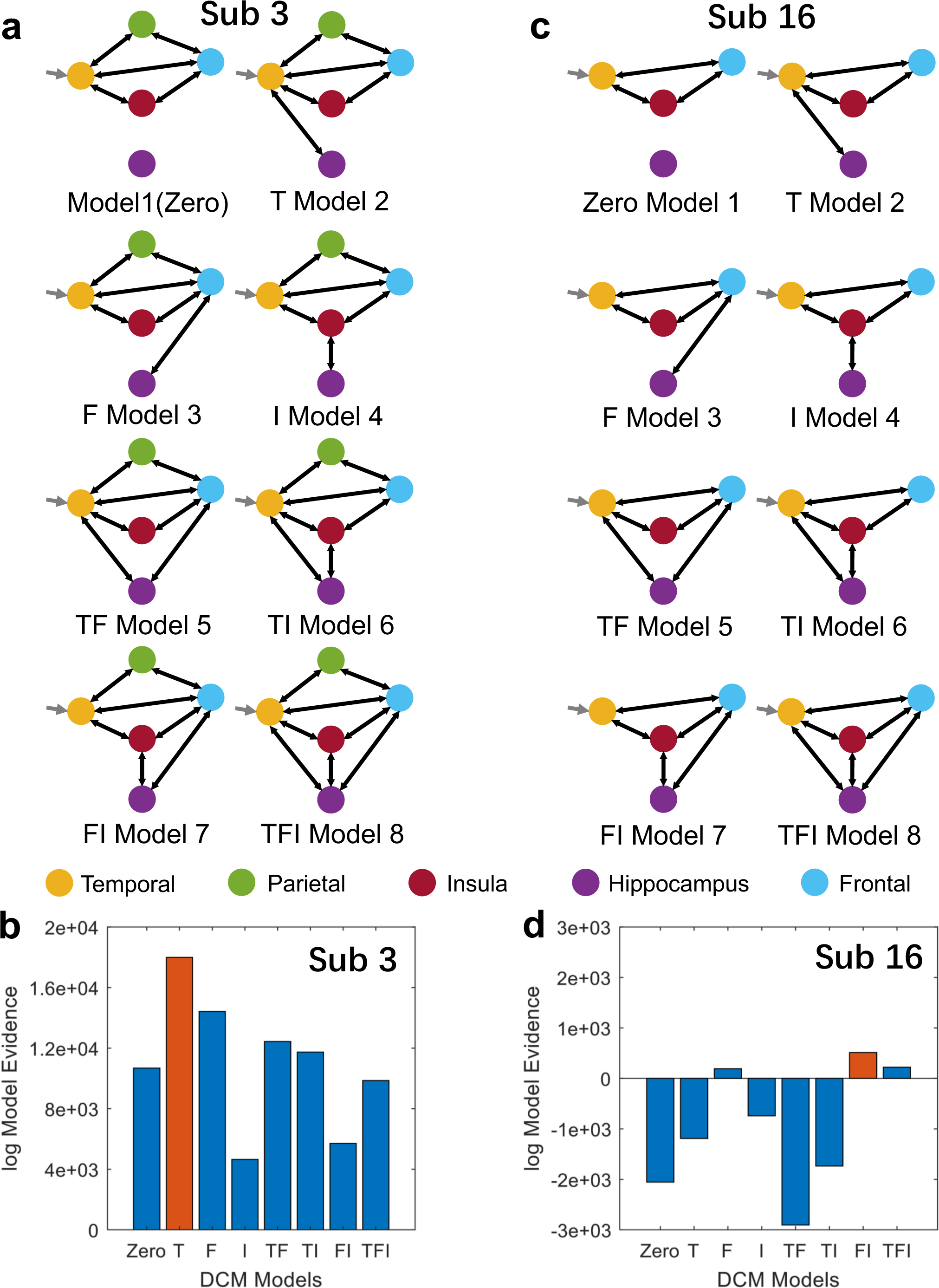

### Intracranial correlates of the mismatch response

Intracranial event-related potentials (iERPs) revealed a robust neural correlate of the auditory mismatch response. Significant deviant-minus-standard difference potential, which is an intracranial counterpart to the scalp MMN^44^, was observed across 188 channels in all 17 patients (Fig. S2). To characterize the dominant response patterns, we performed principal component analysis (PCA) on the iERP waveforms. This analysis identified four distinct spatial-temporal response profiles (Fig. S3). The majority of responses corresponded to a positive-going mismatch response (MMR): Group 1 (121 channels, 6 patients), Group 2 (41 channels, 6 patients), and Group 4 (8 channels, 4 patients) all exhibited a significant positive MMR (p < 0.05). In contrast, a smaller subset of responses in Group 3 (18 channels, 3 patients) showed a significant negative deflection, consistent with the classic mismatch negativity (MMN) pattern (p < 0.05). Given that the same neural phenomenon can be recorded either as a positive or a negative iERP^30^, these results demonstrate that the neural generators underlying deviance detection are broadly accessible in intracranial recordings and exhibit heterogeneous polarity, with a positive MMR being the predominant signature in our dataset.

## Discussion

An ASTM trace constitutes the specific, sustained pattern of neural activity that physically represents and temporarily maintains information, serving as the neurobiological substrate of auditory short-term memory. In the current study, we employed a short-term memory paradigm featuring a contrastive auditory stream of pure tones to investigate the role of the human hippocampus in the formation of auditory short-term memory (ASTM) trace. Combined with the high spatiotemporal resolution approach, intracranial recordings used in this study allowed us to examine the electrophysiological activity across multiple brain regions. Using high frequency activity (HFA, 70–150 Hz) as a reliable correlate of localized neuronal firing, we found that deviance detection elicits significant HFA, indicating ASTM trace generation, and revealed hippocampus is involved in the process. Furthermore, the response latencies (110–438 ms) observed in the hippocampus, insula, temporal, parietal, and the frontal regions suggest that the initial process of the ASTM trace occurs at a pre-attention stage. Functional connectivity obtained from Granger causal analysis based on state-space (SS) model showed that ASTM trace processing engages a hierarchical cortical network, with early bottom-up information flow from the insula and auditory regions to the frontal cortex, followed by later top-down information flow from the frontal cortex to the hippocampus, inferior temporal cortex, and insula. Furthermore, to move beyond this descriptive account and identify the neurobiological mechanism underlying this information flow, we applied dynamic causal modeling. DCM results revealed a convergent neural mechanism: deviant stimuli selectively enhanced the effectiveness of top-down connections directed to the hippocampus, indicating that the hippocampus receives and integrates selectively amplified top-down signals during ASTM trace formation. In summary, these findings provide direct evidence that the human hippocampus contributes to ASTM trace formation at a pre-attention stage via a stimulus enhanced top-down pathway. Our study offers a more comprehensive model of ASTM trace formation and challenges entrenched definition of memory systems.

### The hippocampus is involved in pre-attentive auditory short-term memory trace formation

A conventional view in memory neuroscience posits that the human hippocampus plays a crucial role in long-term, or episodic memory but does not contribute to short-term memory^2^. Consistent with this view, previous research has established that the hippocampus participates in auditory processing, often related to later perceptual and cognitive processes^33–36^. For instance, prior research using electrophysiological techniques revealed that auditory processing engages a distributed network and that the hippocampal contribution to the scalp P300 is spatially restricted to the anterior temporal region^45^. Moreover, hippocampal functional engagement has been characterized in contexts of higher-order cognitive operations, including associative learning like auditory fear conditioning^46^ and contextual updating at the end of a cognitive episode^47^. Anatomically and physiologically, the auditory association cortices send afferent inputs to the hippocampus, which in turn projects back to the primary auditory cortex and auditory association areas during auditory perception^48,49^.

This entrenched view, however, is increasingly difficult to reconcile with findings that place the hippocampus with rapid, pre-attentive processing circuits. Notably, MEG study showed hippocampus forms a functional circuit with the ventromedial prefrontal cortex (vmPFC) during mismatch detection^50^, suggesting its engagement occurs at earlier, more fundamental processing stages than previously theorized. The MMN has been regarded as the index of ASTM trace formation and deviance detection for decades, it is known to reflect a cortical comparison process between deviant stimuli and memory trace of standard stimuli^19–26^. While the scalp-recorded, low-frequency ERP and the low spatial resolution limits its ability to delineate the ASTM trace at the neuronal level. Here, we used sEEG, which provides excellent spatio-temporal resolution and unique access to deep limbic structures like the hippocampus^51^, and measured HFA (70lJ150 Hz) as a more direct index of ASTM trace formation. HFA is a well-established signature of localized cortical firing^31^ that offers greater temporal and spatial specificity than the integrated, scalp-recorded MMN for tracking rapid network dynamics^52^. In this study, we found that auditory deviant stimuli elicit widespread high-frequency activity (HFA: 70–150 Hz) with latencies ranging from 110 to 438 ms, indicating that ASTM trace formation occurs at a pre-attentive stage and that the hippocampus is involved in this process. This finding fundamentally challenges the traditional segregation of memory systems. The neural responses of hippocampus at a pre-attentive stage indicate that it is not exclusive to long-term memory encoding, but is involved in the initial, rapid construction of ASTM trace. We therefore propose that ASTM trace formation is supported by a cortico-hippocampal network. This repositions the hippocampus with a newly defined and foundational role in short-term memory.

### Hierarchical information flow during ASTM trace formation

To clarify the specific mechanism of the hippocampus contributes to ASTM trace formation, we first characterized the spatiotemporal dynamics of information flow using Granger causality (GC) analysis based on a state-space model. Functional connectivity results obtained by GC analysis revealed that ASTM trace processing follows a precise spatiotemporal and hierarchical organization/sequence as follows (Fig. 4c and Fig. 5). An early bottom-up information flows from the insula and auditory-related regions to the frontal lobe, followed by later top-down information flow from the frontal lobe back to the hippocampus, inferior temporal lobe, and insula at the pre-attentive stage. This finding not only aligns with the core predictive coding principle, in which bottom-up signals convey prediction errors and top-down signals convey updated predictions^52–54^, but also precisely positions the hippocampus as an integration node within the hierarchical network of ASTM trace formation.

### The mechanism of hippocampus contributes to auditory short-term memory trace formation

To further uncover the underlying mechanism, we applied Dynamic Causal Modeling (DCM) to analyze effective connectivity. Bayesian model selection showed that deviant stimuli selectively enhanced the effectiveness of top-down connections directed to the hippocampus, including temporal cortex to the hippocampus, the frontal cortex and the insula to the hippocampus (Fig. 6a and Fig. 6b). This indicates that hippocampal engagement in ASTM trace formation is not a mere epiphenomenon, but is governed by a stimulus-specific, top-down gain-control mechanism. The early engagement of the insula aligns with its well-established role in auditory processing. Converging evidence indicates that the insula is crucial for basic auditory detection, multisensory integration, as well as in allocating auditory attention and orienting to novel stimuli^55,56^. As shown by prior iEEG work, the insula is involved in automatic auditory processing^57^, this aligns with our observation of its early engagement in the bottom-up stream. Additionally, the DCM based regulatory network features a direct top-down connection from the insula to the hippocampus. It also incorporates bidirectional regulation between the frontal and temporal cortices and a direct frontal-hippocampal pathway, together forming a multi-nodal, cascaded regulatory architecture. It is also important to note that the parietal region participates in the hierarchical circuit supporting ASTM trace formation, albeit via frontal and temporal relays to the hippocampus rather than through direct connections (Fig. 6a, T model). This finding is consistent with the view that STM has been considered to be the preserve of frontoparietal networks. Thus, we suggest that the hippocampus does not operate in isolation from classic STM system but is organized within an extended fronto-parietal-temporal-hippocampal network. This helps understand the apparent paradox between the hippocampus’s role in long-term memory and its involvement in pre-attentive deviance detection. Hippocampus acts as a hub that is recruited at the early stage of memory formation, binding significant sensory events into ASTM under top-down control.

In summary, by combining the directional and temporal network dynamics revealed by GC analysis with the connection-specific regulatory mechanism identified by DCM, this study provides converging evidence that the human hippocampus contributes to ASTM trace formation during a pre-attentive stage via a stimulus-modulated top-down pathway. These findings challenge the classical dichotomy of memory systems and underscore the hippocampus’s active role in the early construction of short-term memory traces.

## Materials and methods

### Ethics statement

This study was approved by the Research Ethics Committee of the First Affiliated Hospital, University of Science and Technology of China (2024KY271). All participants provided written informed consent prior to taking part in the study. Neither the study procedures nor the analytical methods were preregistered in a time-stamped institutional registry before the research was conducted.

### Participants

Stereoelectroencephalography (sEEG) data were obtained from 17 patients (6 males, 11 females) with drug-resistant epilepsy who were being evaluated as potential candidates for resective surgery. Data collection took place at the First Affiliated Hospital, University of Science and Technology of China. As part of their presurgical assessment, each patient underwent invasive sEEG monitoring. The selection of regions for electrode implantation was determined solely by clinical criteria. All participants were implanted with intracranial depth sEEG electrodes (0.8 mm in diameter, with a 3.5 mm center-to-center interelectrode distance) to help localize the epileptogenic zone. Every participant was right-handed, had normal cognitive function, and normal hearing. Written informed consent was obtained from all subjects prior to enrollment. The experimental protocol received approval from the Institutional Review Board of the University of Science and Technology of China and the First Affiliated Hospital.

### Task paradigm

Stimuli in the current study consisted of pure tones generated using Audition 3.0 (Adobe Systems Inc., Mountain View, CA, USA). The auditory contrast was formed by a sequence of 300 ms pure tones, with a 500 Hz tone serving as the frequent standard stimulus and a 700 Hz tone as the infrequent deviant stimulus (**Figure 1**). All stimuli were delivered in random order via headphones using E-Prime (Psychology Software Tools, Inc., USA). Participants were instructed to ignore the auditory stimuli and instead focus on watching a muted movie of their choice with subtitles.

### Data acquisition

For each participant, structural MRI scans were acquired prior to electrode implantation, and post-implantation CT scans were obtained afterward. sEEG signals were recorded using a 128/256-channel Neurofax EEG-1200c amplifier/digitizer system (Nihon Kohden, Japan) via implanted depth electrodes. Recording settings included a high-pass filter with a 0.01 Hz cut off, a 50 Hz notch filter, and a sampling rate of either 2000 Hz or 5000 Hz. Electrodes located on the inner surface of the skull served as the ground and reference.

### Preprocessing and Electrode localization

Our sEEG data was imported to MATLAB using FieldTrip toolbox^58^ and resampled to 1000 Hz. Event onset markers were read from pre-allocated trigger channels. The data were then re-referenced to the mean of the mastoid electrodes, and unused channels were removed. Segments containing epileptiform activity or abnormal fluctuations were manually rejected. The electrode localization was based on the FieldTrip protocol^59^ with an adaptation for Windows, in which post-implantation CT images was co-registered to pre-implantation MRI images using SPM 12^60^. The images were subsequently normalized to the MNI-152 template space using SPM12^61^, and electrode positions were determined according to the FieldTrip pipeline.

### High frequency activity (HFA) analysis

Line noises at 50 Hz, 100 Hz, and 150 Hz was removed using a notch filter. The continuous data were segmented into 900 ms epochs (lJ200 ms to 700 ms). Trials were rejected if the amplitude exceeded 5 SD standard deviations from the mean for more than 25 consecutive ms, or if the power spectral density exceeded 5 standard deviations for more than 6 consecutive Hz^62^. Eligible trials were re-referenced to a neighboring channel for HFA analysis. HFA signals from –100 ms to 600 ms were extracted for further processing. Preprocessed data were band-pass filtered into eight 10-Hz-wide bands ranging from 70 to 150 Hz using a zero-phase Hamming windowed FIR filters (order 138)^63^. The envelope of each narrowband signal was extracted by taking the absolute value of the analytic signal from the Hilbert transform, and each amplitude time series was divided by its mean value^64^. For channels exhibiting a significant linear trend (R-square > 0.3), the trend was removed. Each channel’s baseline period was removed and normalized by dividing the standard deviants of all trials of the baseline period. Auditory stimulus-evoked neural activity exhibits both enhancements and suppressions, which are spatiotemporally distinct and originate from separate neural mechanisms. Previous studies on auditory cortical responses have consistently demonstrated that the brain displays both increased and decreased activity following sound stimulation, with these opposing responses being generated by different neural processes. It is therefore methodologically necessary, both mathematically and neurobiologically, to model and analyze these responses separately. Notably, stimulus-induced suppression shows characteristic spatiotemporal patterns that are significantly different from those of enhancement, further supporting their generation by distinct neural circuits. Evidence from auditory response research confirms that enhancement and suppression reflect dissociable neurophysiological processes. Thus, independent treatment of these responses is essential for both quantitative and neurobiological interpretations.

### Tone response and deviance detection

To identify channels with significant responses to auditory stimuli, a two-tailed cluster-based permutation test with threshold-free cluster enhancement^37^ and 1000 sign-flipping randomizations^65^ was applied separately to standard and deviant trials for each channel. Channels with a corrected p-value below 0.05 either stimulus type were considered responsive. Channels displaying abnormal waveforms or epileptic activity were subsequently excluded through manual review. Then, in order to assess whether a channel was involved in deviance detection, a two-tailed cluster-based permutation test with threshold-free cluster enhancement^37^ and 1000 randomizations was applied to compare trial categories (standard vs. deviant) for each channel. Channels with a corrected p-value less than 0.05 were considered significant for deviance detection. Additional manual rejection was applied to exclude channels with aberrant signals or association with epileptic activity. The latency of high-frequency activity (HFA) responses was estimated using a fractional area method^66^. The procedure began by identifying the time point of maximum amplitude within the response window as the initial seed point. The significant cluster was expanded bilaterally from this seed until the amplitude dropped below 20% of the seed value or reached the epoch boundaries. The latency was defined as the center of mass of the resulting suprathreshold cluster.

### Event-related potential analysis and clustering

Event-related potentials (ERP) were derived from the same trials used in the high-frequency activity (HFA) analysis, prior to re-referencing to neighboring channels. The ERP signals were filtered with a zero-phase FIR filter (order 18, 1–20Hz), segmented into 700 ms epoches (–100 ms to 600 ms relative to stimulus onset) and baseline-corrected by subtracting the mean amplitude during the –100 ms to 0 ms interval. For channels exhibiting a significant linear trend (R-square > 0.3), detrending was applied. Similar to the HFA analysis, a two-tailed cluster-based permutation test incorporating threshold-free cluster enhancement (TFCE) was conducted for each electrode shaft to identify channels with significant responses to deviant stimuli. Using principal component analysis (PCA) in line with Blenkmann et al.^62^, we identified groups of event-related potential (ERP) responses exhibiting consistent temporal patterns across channels and subjects. This approach also enabled the mitigation of polarity reversal effects. The analysis proceeded as follows: first, a difference waveform was computed by subtracting the standard stimulus response from the deviant stimulus response. PCA was then applied to this difference signal, and the first four components capturing the highest channel weights were retained. Channels were grouped based on their highest component weight, those predominately weighted on the first component were clustered together, followed by those associated with the second component, and so forth for each subsequent component.

### Granger Causality Analysis

Wiener-Granger causality (G-causality, GC) is a statistical notion of causality applicable to time series data, whereby cause precedes, and helps with prediction^67,68^, which has been developed to a powerful method for identifying information transfer or directed functional (“causal”) connectivity in neural time series data^69,70^. It is defined in both time and frequency domains, and allows for the conditioning out of common causal influences. Originally developed in the context of econometric theory, it has since achieved broad application in the neurosciences and beyond^71^. G-causality says that a variable X “G-causes” another variable Y if the past of X contains information that helps predict the future of Y, over and above the information already in the past of Y itself (and in the past of other “conditioning” variables Z). In general terms, when this condition is satisfied one can say that there is “information flow” from X to Y. This is justified because G-causality is an approximation to transfer entropy (the approximation is exact for Gaussian variables), which is a directed version of Shannon’s mutual information, a very general way of characterizing the statistical dependency or shared information between two variables^72^. The multivariate Granger causality analysis (MVGC) Matlab toolbox used in the current study was designed for applications in neuroscience^42^, which implements numerical routines for calculating multivariate Granger causality (MVGC) from time series data, both unconditional and conditional, in the time and frequency domains^43^. Since it has been recognized that a moving average (MA) component in the data presents a serious confound to Granger causal analysis, as routinely performed via autoregressive (AR) modeling^42^. Previous studies have solved this problem by demonstrating that Granger causality may be calculated simply and efficiently from the parameters of a state-space (SS) model^68^. Because SS models are equivalent to autoregressive moving average models, Granger causality estimated in this fashion is not degraded by the presence of a MA component. State-space Granger causal inference stands to significantly enhance our ability to identify and understand causal interactions and information flow in the current study^73^. We used the MVGC (Multivariate Granger Causality) Matlab toolbox for calculating multivariate Granger causality from time series data^42^. Autoregressive moving average (ARMA) process or, equivalently, finite order linear state-space (SS) have been applied in this study^73^. We selected a time window of interest spanning –100 ms to 600 ms to perform identical analyses. The model order was estimated using Akaike information criterion. The significance of difference of G-causality between standard and deviant conditions was estimated using a permutation test.

### Dynamic causal modeling

To test whether the top-down information flow identified by Granger causality analysis originates from direct neural connections that are specifically modulated by deviant stimuli, and to elucidate the underlying neural mechanisms, we conducted Dynamic Causal Modeling (DCM). Participants for DCM analysis were selected based on stereotactic electroencephalography (sEEG) electrode coverage and neural response characteristics. Only participants who exhibited a clear hippocampal mismatch response along with significant activation in multiple other relevant brain regions (e.g., temporal, frontal, and insular cortices) were included. Two participants (Subjects 3 and 16) met all criteria and were analyzed. DCM is a Bayesian framework that employs biophysically constrained neural mass models. Neural sources are modeled as local field potential generators, allowing for inference about the mechanisms underlying evoked responses and quantification of how coupling between sources is modulated by experimental stimuli. Based on prior evidence and the activation patterns observed in this study, network nodes were defined for each participant. For Subject 3, nodes included the hippocampus, insula, and temporal, frontal, and parietal cortices. For Subject 16, nodes included the hippocampus, insula, and temporal and frontal cortices. In line with prior studies suggesting that optimal models of deviance processing incorporate both bottom-up and top-down interactions, our model space included bidirectional connections between the hippocampus and cortical regions. A model space of eight competing models was constructed for each participant. Model 1 served as a baseline (null) model containing no connections to or from the hippocampus^74–76^. Models 2 through 8 were generated by systematically adding bidirectional connections between the hippocampus and the other cortical nodes. Neural activity within a 0–600 ms post-stimulus time window was modeled using a biophysically and spatially constrained local field potential forward model. Finally, Bayesian model selection (BMS) was employed to compare the generative models. BMS evaluates the free-energy bound on the log model evidence, ln p(y|m), which represents the probability of the observed data y under each model m. This metric penalizes model complexity to prevent overfitting. The model with the highest model evidence within the tested set is identified as the “winning” model.

## Data availability

The processed datasets generated and/or analysed in the current study are available from the corresponding author upon reasonable request.

## Authorship contribution statement

X.L., L.C., and R.Q. designed research; X.L. collected sEEG data; R.Q. and X.F. did the electrode implantation surgery for drug-resistant epilepsy patients, Y.Q., D.Z., Y.Z., A.C., L.W., X.L. and W.Z. and provided fMRI and CT files, Z.G. and X.L. analyzed data; X.L. and L.C. wrote the paper.

## Funding

This study was supported by the National Natural Science Foundation of China (Grant GG2070000587) and University of Science and Technology of China (GrantWK3460000008).

## Competing interests

The authors declare that the research was conducted in the absence of any commercial or financial relationships that could be construed as a potential conflict of interest.

---

**Supplementary Figure 1.**
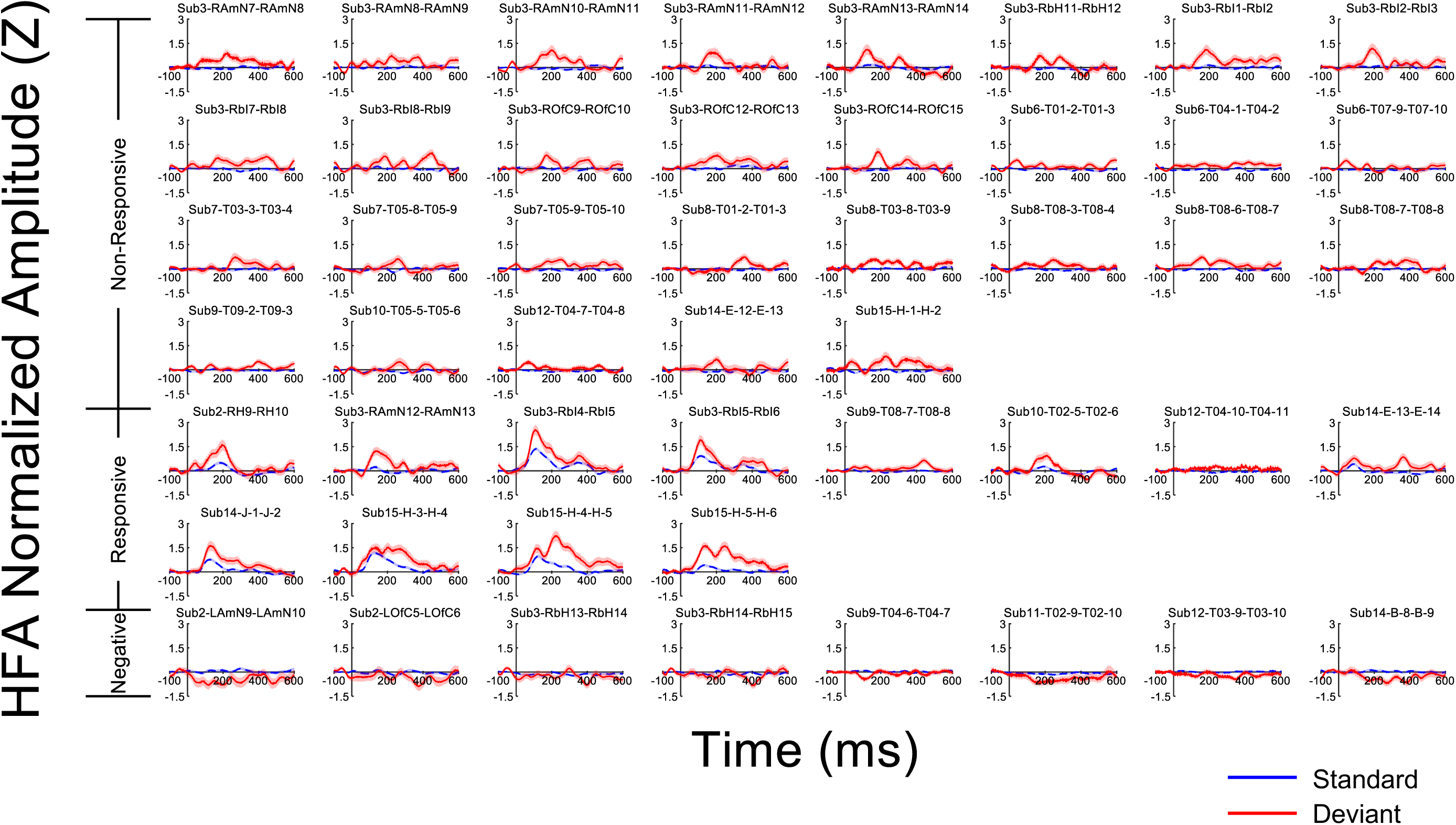

**Supplementary Figure 1.**
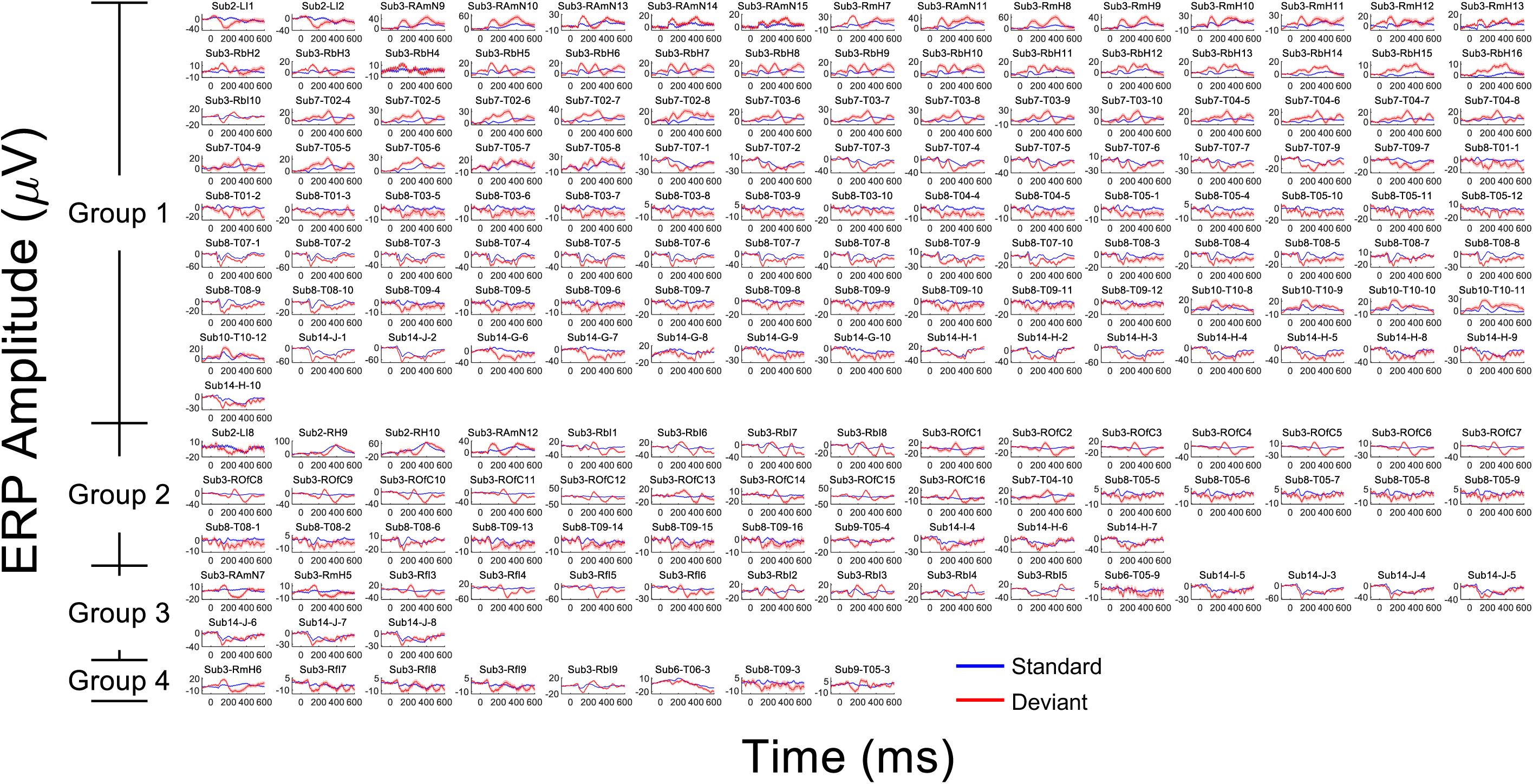

**Supplementary Figure 1.**
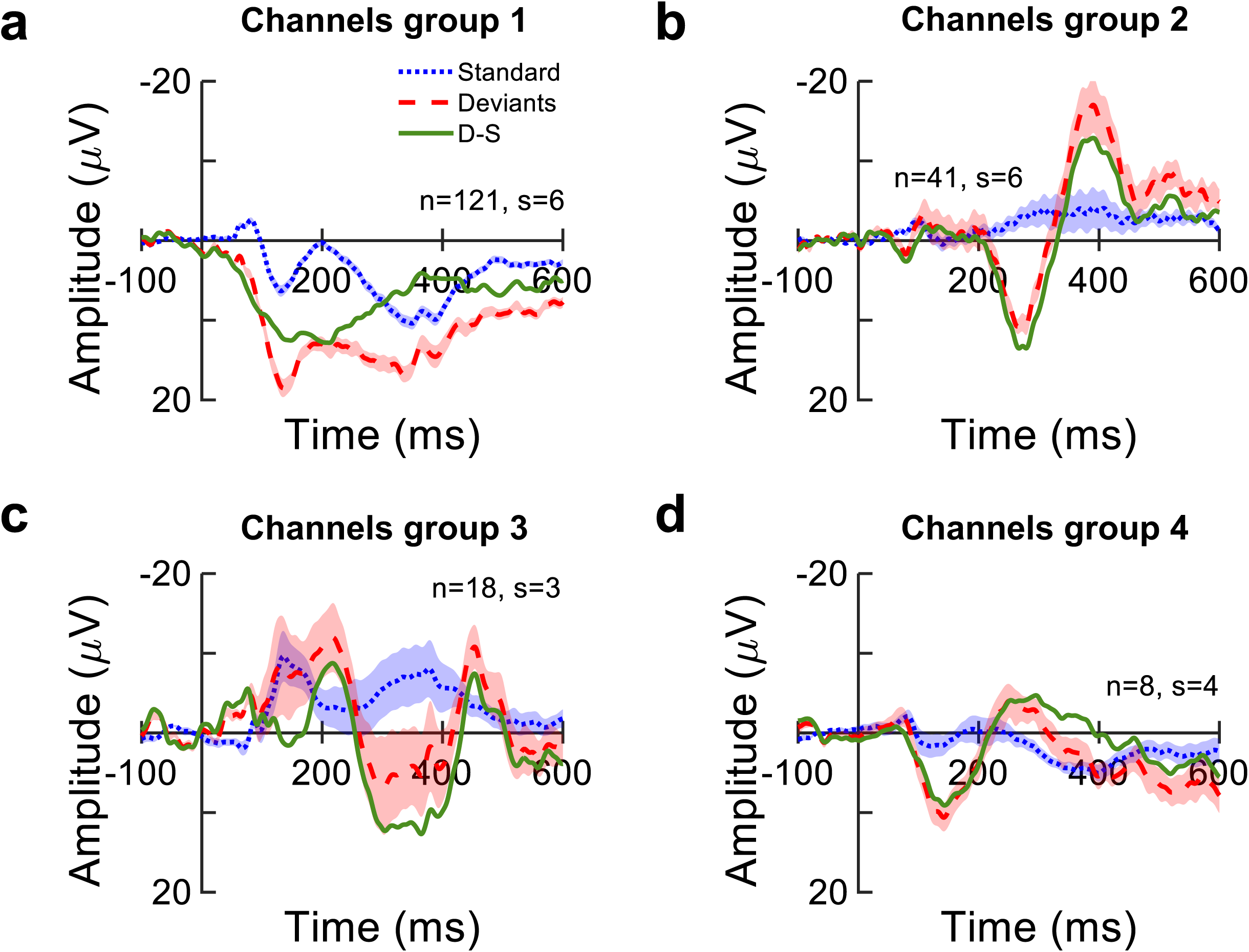

## Notes

### Competing Interest Statement

The authors have declared no competing interest.

### Summary of Updates

Figure 6 as an important new evidence for hippocampus contributes to the auditory short-term memory trace formation. In the main text and methods parts, dynamic causal modeling has been used to analyze the effectivite connectivity of brain regions.

