## Supplementary material for "The human hippocampus is involved in auditory short-term memory trace formation": Table 1

| Patient | Sex (F/M) | Age (Years) | Electrode number | Channel number |
| --- | --- | --- | --- | --- |
| 1 | F | 25 | 6 | 66 |
| 2 | F | 30 | 6 | 60 |
| 3 | F | 28 | 7 | 78 |
| 4 | F | 35 | 10 | 108 |
| 5 | F | 52 | 9 | 86 |
| 6 | M | 32 | 8 | 78 |
| 7 | M | 26 | 10 | 96 |
| 8 | F | 35 | 9 | 96 |
| 9 | F | 33 | 10 | 98 |
| 10 | M | 30 | 10 | 106 |
| 11 | M | 24 | 7 | 70 |
| 12 | M | 35 | 10 | 110 |
| 13 | F | 26 | 12 | 108 |
| 14 | F | 30 | 10 | 114 |
| 15 | F | 34 | 10 | 104 |
| 16 | F | 36 | 8 | 84 |
| 17 | M | 15 | 8 | 108 |

**Table 1. Subjects and channels.**

F: Female; M: Male.
